# In vivo magnetic resonance spectroscopy of hyperpolarized [1-^13^C] pyruvate in a male guinea pig model of life-long Western diet consumption and non-alcoholic fatty liver disease development

**DOI:** 10.1101/2021.02.05.429612

**Authors:** Lauren M. Smith, Conrad B. Pitts, Lanette J. Friesen-Waldner, Neetin H. Prabhu, Katherine E. Mathers, Kevin J. Sinclair, Trevor P. Wade, Timothy R.H. Regnault, Charles A. McKenzie

**Author notes:** Correspondence to: Charles McKenzie, 1151 Richmond Street, Natural Sciences Center 9A, London, Ontario, Canada, N6A 3K7.

## Abstract

**BACKGROUND:** Alterations in glycolysis and oxidative pathways are central to the increasing incidence of non-alcoholic fatty liver disease (NAFLD), highlighting a need for *in vivo*, non-invasive technologies to understand the development of hepatic metabolic aberrations in lean NAFLD.

**PURPOSE/HYPOTHESIS:** To use hyperpolarized magnetic resonance spectroscopy (MRS) and proton density fat fraction (PDFF) MRI techniques to investigate effects of a chronic, life-long exposure to the Western Diet (WD) in a model of NAFLD and identify cellular metabolite changes and correlations related to enzyme activity. It is hypothesized that exposure to the WD will result in NAFLD in association with altered pyruvate metabolism.

**STUDY TYPE:** Prospective POPULATION/SUBJECTS/PHANTOM/SPECIMEN/ANIMAL MODEL: 28 male guinea pigs were weaned onto a control diet or WD.

**FIELD STRENGTH/SEQUENCE:** 3T; T1, T2, IDEAL, broadband PRESS MRS.

**ASSESSMENT:** Median PDFF was calculated in the liver and hind limbs. [1-^13^C]pyruvate dynamic MRS in the liver was quantified by the time to peak (TTP), calculated as the time from pyruvate peak to metabolite peak. After a recovery period, animals were euthanized, and tissue was analyzed for lipid and cholesterol concentration and enzyme level and activity.

**STATISTICAL TESTS:** Unpaired Student’s t-tests were used to determine differences in measurements between the two diet groups. The Pearson correlation coefficient was calculated to determine correlations between measurements.

**RESULTS:** Life-long WD consumption resulted in significantly higher liver PDFF correlated with elevated triglyceride content in the liver. The WD group exhibited a decreased TTP for lactate production, and *ex vivo* analysis highlighted increased liver lactate dehydrogenase (LDH) activity. DATA CONCLUSION: PDFF MRI results suggest differential fat deposition patterns occurring in animals fed a life-long WD, corresponding with increased liver triglyceride levels characteristic of lean NAFLD. The decreased liver lactate TTP and increased *ex vivo* LDH activity suggest lipid accumulation occurs in association with a shift from oxidative metabolism to anaerobic glycolytic metabolism in WD livers.

## Introduction

With increasing consumption of the high fat/high sugar ‘Western diet’ (WD), there has been a corresponding increased incidence of non-alcoholic fatty liver disease (NAFLD) and comorbidities in Western society in both lean and overweight/obese populations (1). Following this trend, NAFLD has become the leading cause of chronic liver disease in developed nations (2), imposing significant burdens on healthcare systems and decreasing overall life expectancy (3). Understanding liver metabolism may provide a means of monitoring NAFLD severity as it has previously been reported that altered pyruvate metabolism is an indicator of liver disease (4). The increased production of lactate from pyruvate has been highlighted as an indication of the shift from oxidative metabolism to anaerobic glycolysis (5) and is associated with liver damage and also highlighted in other conditions including cancer, heart failure, and neurodegeneration (4, 6). Elevated lactate in the liver, induced by a high fat diet, has been found in obese mice (7) and may be linked to a disturbance in hepatic lipid synthesis (6). Increased lactate is associated with diabetes and, specifically, the interference of hepatic glycogen synthesis and insulin dysregulation in the liver (8). The ability to measure lactate production *in vivo* is useful for observing NAFLD’s effects on liver damage, stress, and lipid accumulation.

Magnetic resonance imaging (MRI) is a useful tool that can be used to evaluate structure and function, making it an ideal method for characterizing liver disease. Chemical shift imaging is an MRI technique used to separate signal from fat and water within the body, enabling *in vivo* measurements of fat fractions in any body region. Additionally, MRI can detect signal from nonproton nuclei, allowing for the selective imaging of certain biomolecules of interest. Carbon-13 (^13^C) magnetic resonance spectroscopy (MRS) has historically been used to investigate glycolysis but is limited by inherently low sensitivity and long acquisition times (9). To address these limitations, hyperpolarized ^13^C MRS is used to temporarily boost the signal-to-noise ratio, allowing for rapid acquisition of spectroscopy signal from ^13^C-enriched substrates.

By enriching pyruvate with ^13^C and using hyperpolarized technology to significantly enhance the magnetic resonance signal, it is possible to inject and subsequently image the distribution of [1-^13^C]pyruvate *in vivo* and in real-time. Additionally, this technique allows us to simultaneously acquire and subsequently differentiate signals from the pyruvate molecule and its downstream metabolites that retain the ^13^C nuclei over the acquisition duration. Concentrations and time curves of each metabolite can be quantified, allowing examination of metabolism. Calculated time-to-peak (TTP) is a quantitative indirect measurement of enzyme concentration as the temporal dynamics of the metabolic reactions are directly related to its concentration (10). Animal studies are crucial in understanding the fundamental biochemical properties of disease and validating emerging technologies such as hyperpolarized MRI by correlating ^13^C exchange rates with *ex vivo* measurements of hepatic enzyme activities. Hyperpolarized [1-^13^C]pyruvate MRS has previously been used in a NAFLD rat model where both [1-^13^C]alanine and [1-^13^C]lactate were identified as potentially useful non-invasive markers of the progression of NAFLD.

The current study’s goal was to validate MRI techniques to investigate the effect of long-term WD consumption on liver metabolism in a pre-clinical guinea pig model of NAFLD. The pyruvate metabolic pathway was targeted with hyperpolarized ^13^C MRS to identify pyruvate metabolism changes as potential biomarkers of liver damage. Liver tissue assays were conducted to complement imaging data. This validation will allow us to more confidently utilize these techniques going forward to investigate the severity of metabolic states *in vivo*, and such capability may facilitate the longitudinal assessment of therapeutic response for NAFLD.

## Materials and Methods

### Ethical approval

Animal care, maintenance, and procedures were performed following the national council’s standards and guidelines on animal care. All procedures were reviewed, approved, and monitored by the institutional animal care and ethics committee.

### Animal model and welfare

Guinea pigs were used in this study as they differ from other rodents as a model for NAFLD in that their lipoprotein metabolism and hepatic enzyme activity closely mimics human physiology (11). Male guinea pig pups were born in-house to mothers fed a standard diet throughout gestation and lactation in a 12/12 hour light-dark schedule in individual cages. At approximately fifteen days postnatal (PN), pups were weaned onto their respective diets, feeding *ad libidum* in individual cages. Male guinea pig pups (matched for litter) were randomly weaned onto either a control diet (CD: 21.6% protein, 18.4% fat, 60% carbohydrates, n = 15) or WD (21.4% protein, 45.3% fat, 33.3% carbohydrates, n = 14) (12). Percentages indicate the calorie contribution from each macronutrient to the total dietary calories. The fat content (CD: 3% saturated fatty acids (SFA), 4% monounsaturated fatty acids (MUFA), 11% polyunsaturated fatty acids (PUFA); WD: 32% SFA, 12% MUFA, 2% PUFA) and carbohydrate content (CD: 10% sucrose, 40% corn starch; WD: 19% sucrose, 6.5% fructose, 9% corn starch; % by weight) of the diets differed in terms of their compositions. The WD had a higher caloric density (4.2 vs 3.8 kcal/g) and included 0.25% cholesterol (12). Daily food consumption (g/day/kg body weight) and animal weights were recorded for the 10 days before MRI scanning and during the period between MRI and euthanisation. At 144 days PN, animals underwent scanning (details below), and at approximately 150 days PN, animals were euthanised by CO_2_ inhalation in a sealed chamber (13). Blood samples were immediately collected from the descending vena cava and analyzed using VetScan VS2 Chemistry Analyzer (VetScan® Mammalian Liver Profile reagent, Abaxis, Union City, CA) to quantify levels of alkaline phosphatase (ALP), alanine aminotransferase (ALT), gamma glutamyl-transferase (GGT), blood ammonia (BA), bilirubin (TBIL), albumin (ALB), blood urea nitrogen (BUN), and cholesterol (CHOL). Livers were then harvested, weighed, snap-frozen in liquid nitrogen, and stored at −80°C until later biochemical determinations.

### In Vivo Proton MRI Determination of Fat Content and In Vivo Measurements of Hepatic Metabolism With Hyperpolarized ^13^C MRS

After 144 +/- 4 days (equivalent to ~ 18-22 human years (14)), animals were imaged using a 3T GE MRI under anesthetic (15). Animals were anesthetized using 4.5% isoflurane with 2L/min O_2_ and maintained between 1.5-2.5% isoflurane with 2L/min O_2_. A catheter was inserted into the hind foot saphenous vein for intravenous administration of the hyperpolarized ^13^C pyruvate during the MRI exam. Vital signs were monitored throughout the experiment, and body temperature was maintained at 37°C. To standardize their metabolic state at the start of the experiment, all animals underwent MRI at roughly the same time of day, and all animals were fasted for 2 hours before imaging and a subcutaneous injection of glycopyrrolate (0.01mg/Kg body weight) was administered half an hour before administration of anesthetic.

Anatomical T1-weighted gradient echo (repetition time/echo time [TR/TE] = 5.1/2.4 ms, flip angle = 15°, number of averages = 4, slice thickness = 0.9 mm, total scan time ~ 7 min) and T2-weighted spin-echo (TR/TE = 2000/120 ms, number of averages = 2, slice thickness = 0.9 mm, total scan time ~ 7min) images with 0.875 × 0.875 mm^2^ in-plane resolutions were obtained using a 32-element cardiac coil (In Vivo Corp., Gainesville, FL). Water-fat images were acquired using a modified IDEAL acquisition (TR/ΔTE = 9.4/0.974 ms, echoes = 6, flip angle = 4°, number of averages = 4, slice thickness = 0.9 mm, total scan time ~ 13 min) with a 0.933 × 0.933 mm^2^ inplane resolution and reconstructed into PDFF images. CSE-MRI (IDEAL-IQ) used parallel MRI to accelerate by a factor of 1.75 in the phase and slice directions. Regions of interest were drawn around the liver, hind limb tissue, whole body, subcutaneous adipose tissue (SAT), and visceral adipose tissue (VAT). These segmentations were done manually with digitizing monitors using 3D Slicer (version 4.10.0, www.slicer.org). For this study, the VAT was defined as the adipose tissue visible on IDEAL fat images below the diaphragm and above the pelvis external to the abdominal cavity organs. SAT and VAT were reported as a percentage of the total body volume, calculated by dividing the adipose tissue volume by the whole body volume. These segmentations were used to calculate each region’s total volume and the median PDFF of each region when overlaid on IDEAL fat fraction images. Areas of fat-water swaps were excluded from volumes that were used to measure PDFF.

Anatomical images were used as a reference to select a slab through the liver for ^13^C MRS. PRESS chemical shift MRS (TR/TE = 1082/35 ms, echoes = 1, slice thickness = 20.4 mm) was used to acquire hyperpolarized ^13^C spectra over 90 s with a 1 s time resolution using a custom ^13^C birdcage coil (Morris Instruments, Ottawa, Canada). [1-^13^C]pyruvate (Cambridge Isotope Labs, Massachusetts, USA) with 15mM Ox063 (Oxford Instruments, Oxford, UK)and 1.5mM Dotarem (Guebert, Villepinte, France) was hyperpolarized (Hypersense, Oxford Instruments), and a 3.5 mL bolus of the 80 mM solution (pH balanced, 37 °C)was injected over approximately 12 s into a vein in the hind leg. Spectra were analyzed using SAGE software (GE Medical Systems), and the time to peak (TTP) was measured as the time between the pyruvate peak and metabolite peak to mitigate effects due to slight differences in injection times. TTP is a model-free analysis metric that roughly displays an inverse correlation with enzyme concentration (10). The animals were monitored, warmed, and kept on 100% O_2_ until they began to wake up. They were then placed under a heating lamp and monitored until they were fully awake and mobile, at which time they were returned to their cages.

### Ex Vivo Hepatic Determinations

#### Triglyceride Content

The left liver lobe was ground into a frozen powder over liquid nitrogen and analyzed for liver triglyceride levels using a colorimetric assay (Cayman Chemicals, Ann Arbor, MI, USA) following the manufacturer’s instructions. Briefly, approximately 200 mg tissue was homogenized in NP-40 buffer containing leupeptin using an electric homogenizer. Samples were centrifuged at 10 000 *g*, and the supernatant was harvested. Samples from CD were not diluted, whereas samples from WD were diluted 1:4 in NP-40 before assaying. After incubating the samples in the enzyme mixture for 15 minutes in the dark, the plate was read at 530 nm, 540 nm, and 550 nm. The absorbance at the three wavelengths was averaged and used to calculate triglyceride concentration based on the standard curve. Triglyceride concentration was normalized to protein concentration by Pierce BCA assay (ThermoFisher, Waltham, MA, USA).

#### Liver cholesterol content

Total lipids were extracted from approximately 150 mg of frozen liver tissue following the Folch method (16). Total cholesterol, free cholesterol, and cholesteryl ester levels in lipid extracts were determined by enzymatic, colorimetric assays (Wako Diagnostics, Richmond, VA, USA) (17) performed through the Metabolic Phenotyping Laboratory in Robarts Research Institute (London, Ontario, Canada). Cholesterol levels were normalized to the mass of the extracted tissue.

#### Western Blot

Protein was isolated from approximately 100 mg of frozen, ground liver tissue using RIPA buffer containing Aprotinin, Leupeptin, PMSF, NaF, and Sodium Orthovanadate (New England BioLabs, Ipswich, MA, USA). Tissue was homogenized using an electric homogenizer, sonicated at 30% amplitude for 5 second processing time, and centrifuged at 12 000 *g* for 30 mins at 4°C. The supernatant was harvested and stored at −80°C until use. Protein samples were prepared in laemmli buffer containing b-mercaptoethanol at a final concentration of 5%. Twenty mg of protein was run through 10% polyacrylamide tris-glycine gels and transferred onto PVDF membrane. Membranes were probed for PDH, phosphorylated PDH (pPDH), and LDH (Table 1) overnight at 4°C. Anti-rabbit secondary antibodies (Table 1), conjugated to HRP, were used to detect primary antibodies by incubating for one hour at room temperature. Proteins were detected using Amersham ECL reagent (GE Healthcare, Chicago, IL, USA) and ChemiDoc imager (BioRad) with ImageLab software. Protein expression was normalized to total protein by amido black staining.

**Table 1.**
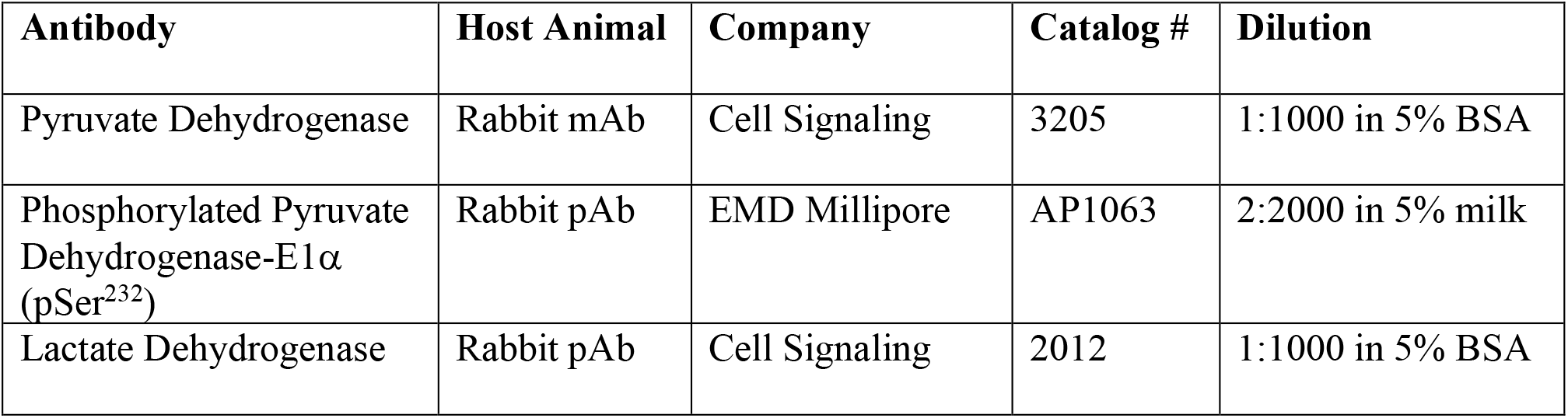
Antibodies used in Western Blot analysis

#### Enzyme Activity Assays

Six samples were randomly selected from each diet group for enzyme activity assays. Liver tissue was homogenized in 9 volumes of homogenization buffer (25 mM HEPES, 2 mM EDTA, 0.1% (v/v) Triton X-100, 10 mM sodium fluoride, 1 mM sodium orthovanadate, pH 7.8) using a microtube plastic pestle. Following incubation at 4°C for 20 min, homogenates were centrifuged at 10000 *g* for 10 min at 4 °C. Floating lipid was removed by aspiration, and pellets were resuspended within the supernatant for each sample. Samples were subjected to three rounds of freeze-thaw in liquid nitrogen, then assayed immediately for enzyme activity. Enzyme assays were performed at 37°C using a Spectramax plate spectrophotometer (Molecular Devices, Sunnyvale, CA, USA) in a 96-well plate. Enzyme activities were expressed relative to total protein concentration for each homogenate (determined by BCA assay).

Lactate dehydrogenase (LDH) activity was measured following the addition of 5ml liver homogenate (diluted 1:20 in homogenization buffer) to 295ml assay mixture containing 50 mM HEPES (pH 7.4), 0.2 mM NADH, 1.0 mM pyruvate. Absorbance values (340 nm) were collected for 3-5 mins, and LDH activity was calculated using an extinction coefficient of 6.22 L mol^-1^ cm^-1^.

Pyruvate dehydrogenase (PDH) activity was measured following the reduction of iodonitrotetrazolium chloride (INT) at 500 nm. Background rates were collected following the addition of 5 ml liver homogenate to 290 ml assay mixture containing 50 mM Tris (pH 7.8), 0.5 mM EDTA, 2.5 mM NAD^+^, 0.2 mM Coenzyme A, 0.1 mM sodium oxalate, 0.4 mM thiamine pyrophosphate, 1 mg/ml bovine serum albumin, 0.1% (v/v) Triton X-100, and 1U/ml diaphorase. The PDH reaction was then initiated by the addition of 5 ml 0.6 M sodium pyruvate. PDH activity was calculated from the difference between the rates with and without pyruvate, using an extinction coefficient of 15.4 L mol^-1^ cm^-1^.

Citrate synthase (CS) activity was measured following the addition of 10 ml liver homogenate (diluted 1:20 in homogenization buffer) to 287 ml assay mixture containing 50 mM Tris (pH 8.0), 0.1 mM 5,5-dithiobis(2-nitro-benzoic acid) DTNB), and 1.15 mM acetyl CoA. Parallel reactions were run with and without the addition of 0.5 mM oxaloacetate. Absorbance values (412 nm) were collected for 5 min, with CS activity calculated from the difference between the rates with and without oxaloacetate, using an extinction coefficient of 13.6 L mol^-1^ cm^-1^.

### Statistical analysis

Unpaired two-tailed Student’s t-tests were used to determine differences in all measurements between animals in the two diet groups. The Pearson correlation coefficient was calculated for correlations between MRI and *ex vivo* data using a two-tailed P-value and a 95% confidence interval. Results are shown as mean ± SEM, and statistical significance was set at p < 0.05. Data analysis was performed using GraphPad Prism 6 (San Diego, CA, USA).

## Results

### Animal Body Weights and Food Intake

Average daily food consumption and daily calorie consumption were not found to significantly differ between the diet groups in the 10 days before MRI (CD 45.27 ± 2.64 g/day/kg body weight vs WD 45.56 ± 4.67 g/day/kg body weight, p = 0.9575; CD 121.1 ± 9.48 kcal/day vs WD 130.6 ± 25.87 kcal/day, p = 0.5784) or during the time between MRI and euthanasia (CD 51.11 ± 2.84 g/day/kg body weight vs WD 54.43 ± 7.58 g/day/kg body weight, p = 0.6671; CD 132.1 ± 11.42 kcal/day vs WD 150.0 ± 33.54 kcal/day, p = 0.4164). On average, guinea pigs in the WD group were found to weigh significantly less than animals in the CD group based on weight recording in the 10 days before (WD 693.6 ± 18.52 g vs CD 770.6 ± 17.97 g, p = 0.0061) and the period after the MRI examination (WD 708.3 ± 19.01 g vs CD 774.6 ± 21.63 g, p = 0.0308).

### Blood profiles show elevated indicators of liver damage in Western Diet animals

ALT levels were significantly elevated in WD animals compared to CD animals (p = 0.0071, Table 2). Blood cholesterol levels were also significantly greater in WD animals compared to CD animals (p < 0.0001, Table 2). No significant differences were observed in levels of ALP, GGT, BA, TBIL, ALB, or BUN (Table 2).

**Table 2.** Liver blood and tissue profiles.

| **Liver Blood Profile** | **<u>CD</u>** | **<u>WD</u>** | **<u>P-value</u>** |
| --- | --- | --- | --- |
| Alkaline phosphate (ALP) [u/L] | 62.43 ± 11.06 | 43.14 ± 5.65 | 0.1301 |
| Alanine aminotranferase (ALT) [u/L] | 48.14 ± 2.80 | 82.00 ± 9.58 | 0.0071 |
| Gamma glutamyl-transferase (GGT) [u/L] | 6.14 ± 0.74 | 6.63 ± 1.53 | 0.7814 |
| Blood ammonia (BA) [μmol/L] | 52.57 ± 13.65 | 65.38 ± 15.14 | 0.5457 |
| Bilirubin (TBIL) [mg/dl] | 0.20 ± 0 | 0.24 ± 0.02 | 0.2914 |
| Albumin (ALB) [g/dl] | 4.03 ± 0.08 | 4.29 ± 0.09 | 0.0508 |
| Blood urea nitrogen (BUN) [mg/dl] | 22.75 ± 1.87 | 23.50 ± 0.98 | 0.7276 |
| Cholesterol (CHOL) [mg/dl] | 71.57 ± 14.37 | 440.4 ± 29.84 | < 0.0001 |
| **Liver Tissue Component** | **<u>CD</u>** | **<u>WD</u>** | **<u>P-value</u>** |
| Triglycerides (TG) [mg/mg protein] | 0.02 ± 0.00 | 0.11 ± 0.01 | < 0.0001 |
| Total cholesterol [mg/mg tissue] | 2.44 ± 0.12 | 19.34 ± 1.91 | < 0.0001 |
| Free cholesterol [mg/mg tissue] | 1.85 ± 0.11 | 4.72 ± 0.28 | < 0.0001 |
| Cholesterol ester [mg/mg tissue] | 0.59 ± 0.07 | 14.62 ± 1.66 | < 0.0001 |

### Impact of life-long Western diet on body and organ fat content

Using MRI, at 144 ± 4 days WD animals had a significantly lower body volume compared to CD animals (WD 631200 ± 16100 mm^2^ vs CD 684800 ± 20000 mm^2^, p = 0.0485; Figure 1A). Additionally, following life-long WD feeding, guinea pigs had significantly elevated liver volume (WD 41400 ± 3200 mm^2^ vs CD 28300 ± 1000 mm^2^, p < 0.0001; Figure 1B) compared to CD animals with no significant differences in hind limb volume (Figure 1C) between the two groups. WD animals showed a significantly elevated liver PDFF (WD 10.64 ± 0.87 % vs CD 6.20 ± 0.34 %, p < 0.0001; Figure 1D) and significantly decreased hind limb PDFF (WD 3.86 ± 0.25 % vs CD 5.47 ± 0.58 %, p = 0.02; Figure 1E) compared to CD animals. While the SAT and VAT as a percentage of total body volume did not show any differences between animals in the two diet groups (Figure 2A, B), there was a reduction in total adipose tissue (as a percentage of total body volume) in WD animals vs CD animals (WD 8.01 ± 0.44 % vs CD 9.32 ± 0.43 %; p = 0.0434; Figure 2C).

**Figure 1:**
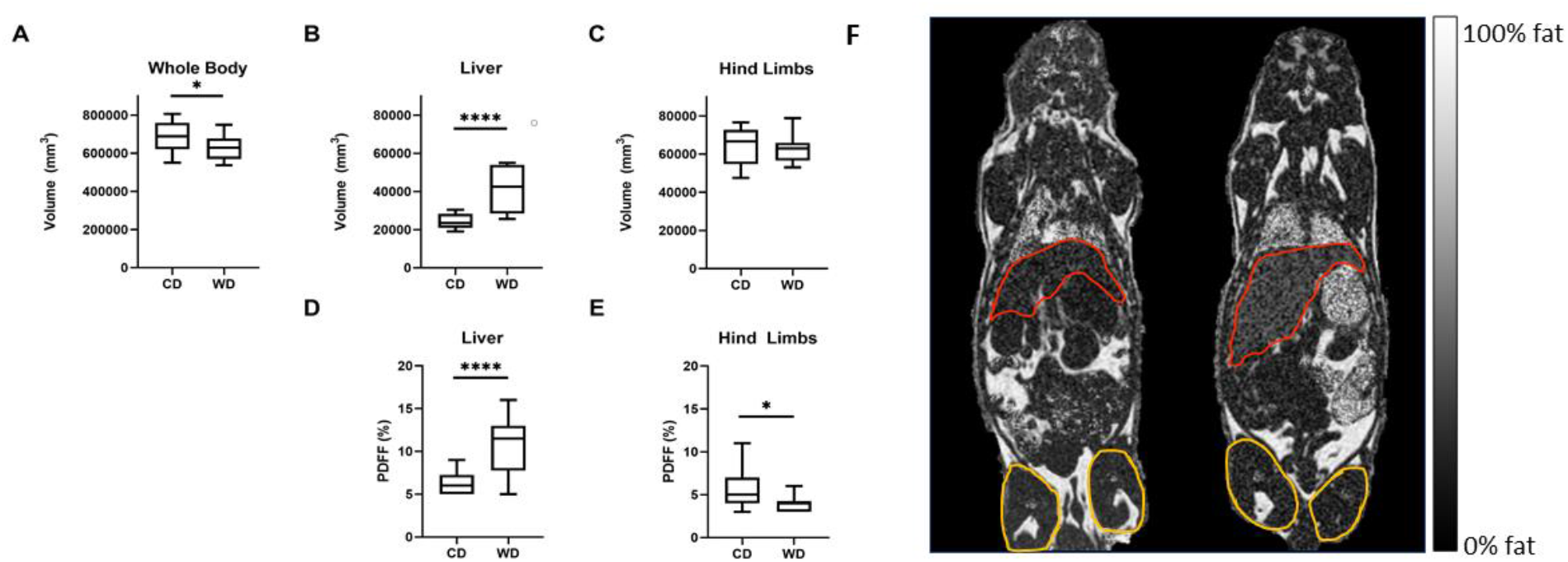
Total volumes estimated from MRI for the whole body (A), liver (B), and hind limbs (C). WD animals displayed a significantly decreased whole-body volume and increased liver volume compared to the CD animals. Median PDFF percentages of the liver (D) and hind limbs (E) show a significantly elevated PDFF in the livers and lower PDFF in the hind limbs of WD animals compared to CD. * indicates p < 0.05, **** indicates p < 0.0001. (F) Examples of proton-density fat fraction image slices from an animal in the CD (left) and WD (right) groups. The livers and hind limbs are outlined in red and yellow, respectively, in both images. The PDFF is visibly elevated in the WD liver, as indicated by a lighter colour, and visibly reduced in the WD hind limb.

**Figure 2:**
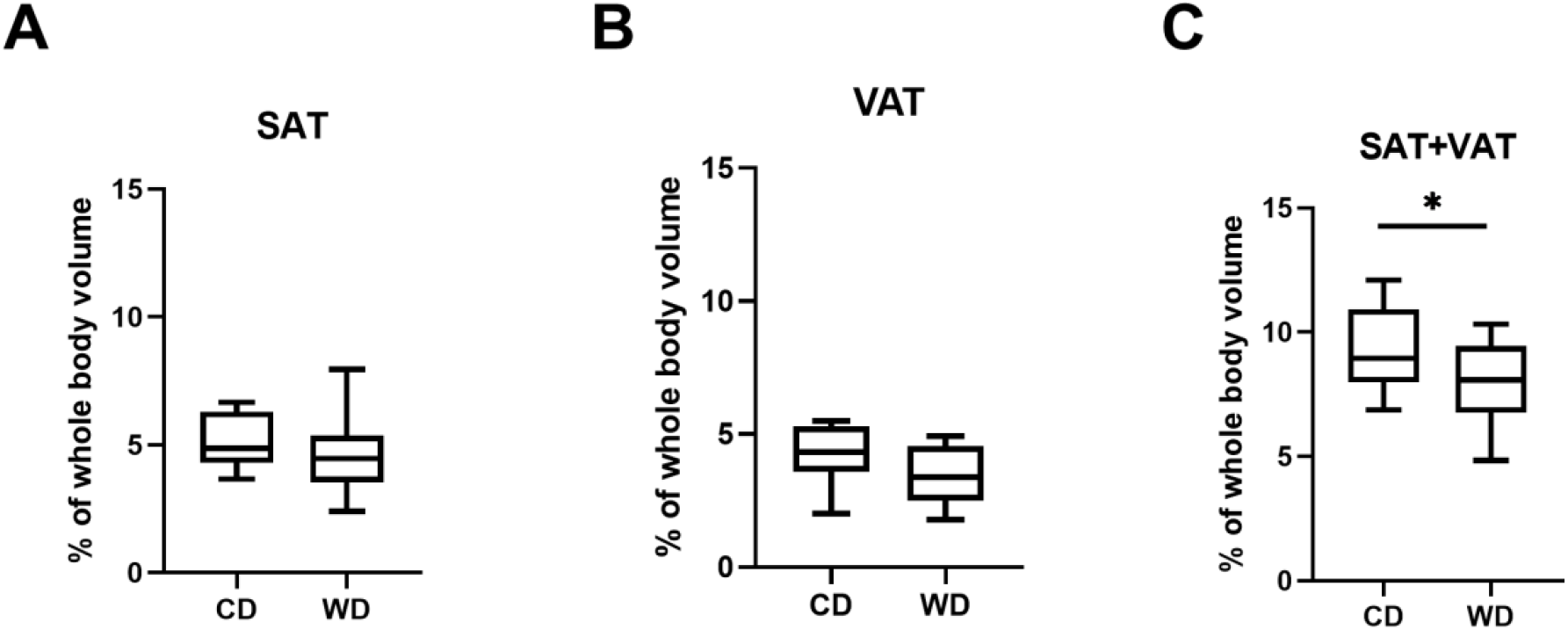
Adipose tissue volumes as a percentage of total body volume for the subcutaneous adipose tissue (A), visceral adipose tissue (B), and the sum of subcutaneous and visceral adipose tissue (C). The SAT+VAT as a percentage of total body volume was significantly elevated in CD animals. * indicates p < 0.05.

### Western diet feeding results in accelerated hepatic lactate production rate

Of the 28 animals scanned in MRS experiments, 26 spectra produced viable data (CD n = 13, WD n = 13; Figure 3A). Lactate TTP in WD animals was significantly lower than in CD animals (WD 214.92 ± 0.75 sec vs CD 11.15 ± 0.54 sec; p = 0.0234; Figure 3B). The TTP related to the rate of metabolism for pyruvate to alanine did not significantly differ between feeding conditions (Figure 3C).

**Figure 3:**
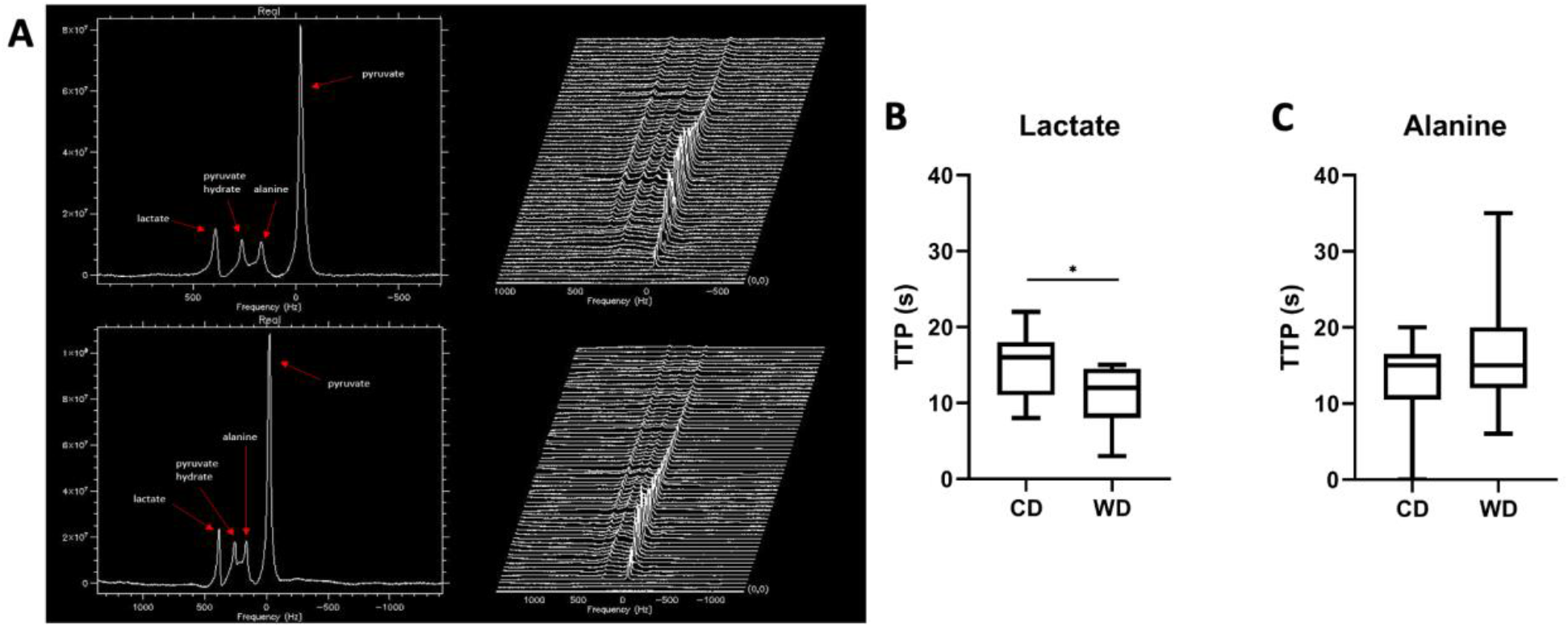
(A) Examples of hyperpolarized [1-^13^C]pyruvate magnetic resonance spectra (left) and stack plots (right) from one CD (top) and one WD (bottom) guinea pig liver. Frequency is relative to the center of the pyruvate peak. Stack plots display spectra from the first 60 seconds of acquisition with a 1 second time resolution. Mean time to peak (TTP) measured from the time of the pyruvate peak for lactate (B) and alanine (C) in both diet groups. WD animals show a significant decrease in lactate TTP compared to CD animals.* indicates p < 0.05.

### Life-long Western diet alters body and liver weights

At 150 ± 6 days of age, tissue collection samples showed WD guinea pigs were significantly lighter than CD guinea pigs (WD 718.8 ± 17.9 g, n = 14 vs CD 787.6 ± 21 g, n = 14; p = 0.0194), despite WD livers being significantly heavier than CD livers both in absolute weight (WD 42.33 ± 2.47 g vs CD 26.70 ± 1.18 g; p < 0.0001) and as a fraction of total body weight (WD 0.059 ± 0.003 vs CD 0.034 ± 0.001; p < 0.0001). Kidneys from WD animals were found to be lighter than those from CD animals as an absolute weight (WD 5.00 ± 0.12 vs CD 5.45 ± 0.16 g; p = 0.0411) but no differences were found in kidney weight as a fraction of total body weight (WD 0.0070 ± 0.0002 vs CD 0.0070 ± 0.0002; p = 0.8828). Brain and heart weights were not significantly different between the two diet groups in absolute weight (WD 3.91 ± 0.24 g vs CD 3.96 ± 0.31 g, p = 0.55; WD 2.84 ± 0.58 g vs CD 3.28 ± 0.56 g, p = 0.15) or as a fraction of total body weight (WD 0.0051 ± 0.0002 vs CD 0.0055 ± 0.0001, p = 0.0721; WD 0.0039 ± 0.0005 vs CD 0.0042 ± 0.0005, p = 0.3226).

### Triglyceride and cholesterol levels are elevated in WD livers

WD animals had significantly elevated hepatic triglycerides compared to CD animals (p < 0.0001, Table 2). The hepatic triglyceride concentration displayed a moderate correlation to the PDFF in the livers of animals in both diet groups (r = 0.6917, Figure 4). Total cholesterol (p < 0.0001; Table 2), free cholesterol (p < 0.0001; Table 2), and cholesteryl ester (p < 0.0001; Table 2) were significantly increased in the WD liver tissues

**Figure 4:**
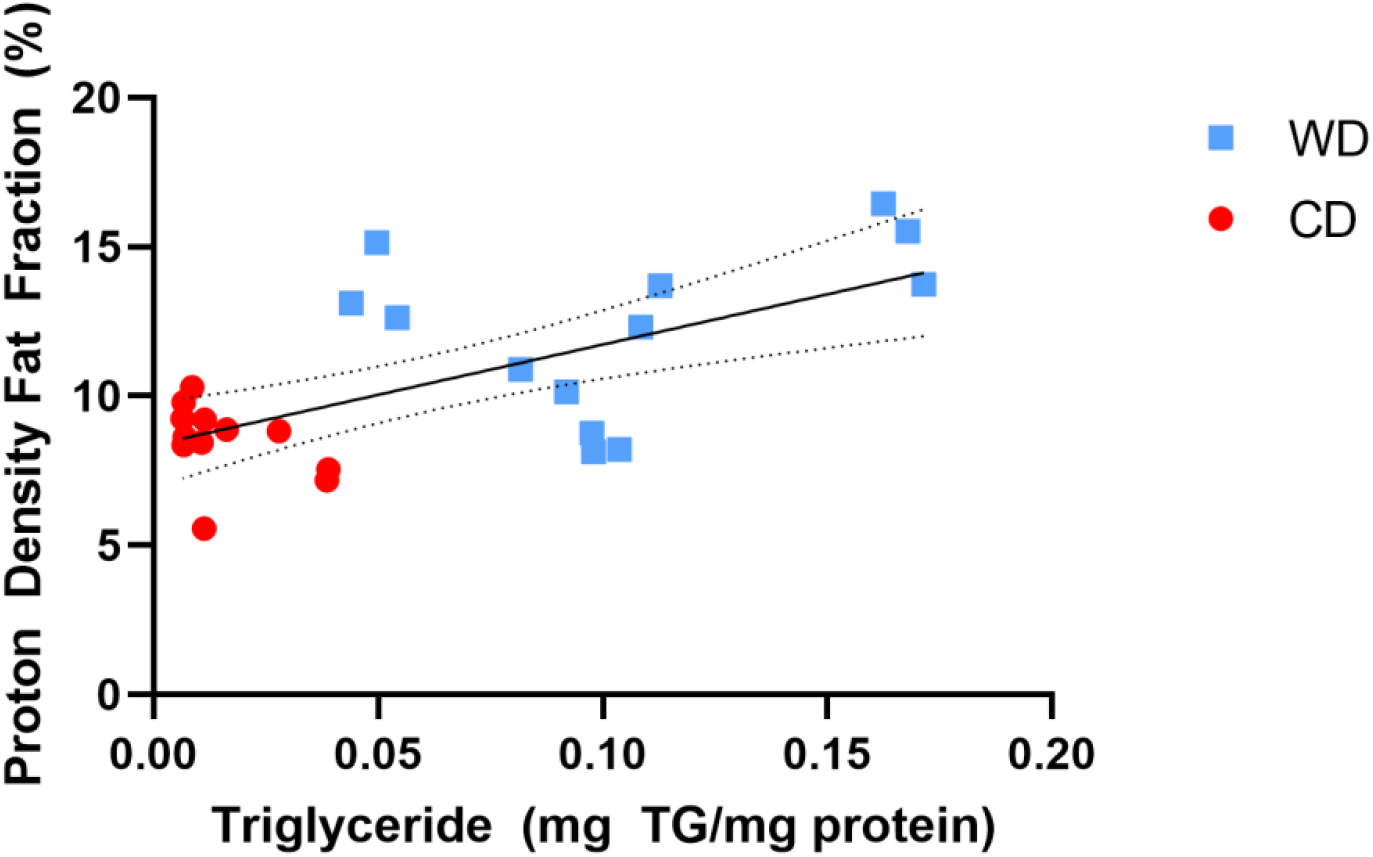
Liver triglyceride concentrations plotted against PDFF in the liver. There is a moderate positive correlation (r = 0.6917, p = 0.0001) between TG and PDFF for all animals. The linear fit line is shown with 95% confidence intervals.

### Altered PDH and LDH activity in WD livers

PDH protein levels were significantly increased in WD animals compared to CD animals (WD 0.42 ± 0.04, n = 14 vs CD 0.28 ± 0.02, n = 13; p = 0.0065, Figure 5A), with no difference observed in phosphorylated PDH (Figure 5B), although PDH activity was significantly decreased in WD animals (WD 0.50 ± 0.24 μmol/min*mg protein, n = 5 vs CD 1.56 ± 0.14 μmol/min*mg protein, n = 6; p = 0.0034, Figure 5D). Despite no difference in protein levels of LDH (Figure5C), LDH activity was significantly elevated in WD (WD 638.6 ± 73.1 μmol/min*mg protein, n = 6 vs CD 388.7 ± 35.3 μmol/min*mg protein, n = 6; p = 0.0116, Figure 5E). No significant difference was observed in CS activity (Figure 5F).

**Figure 5:**
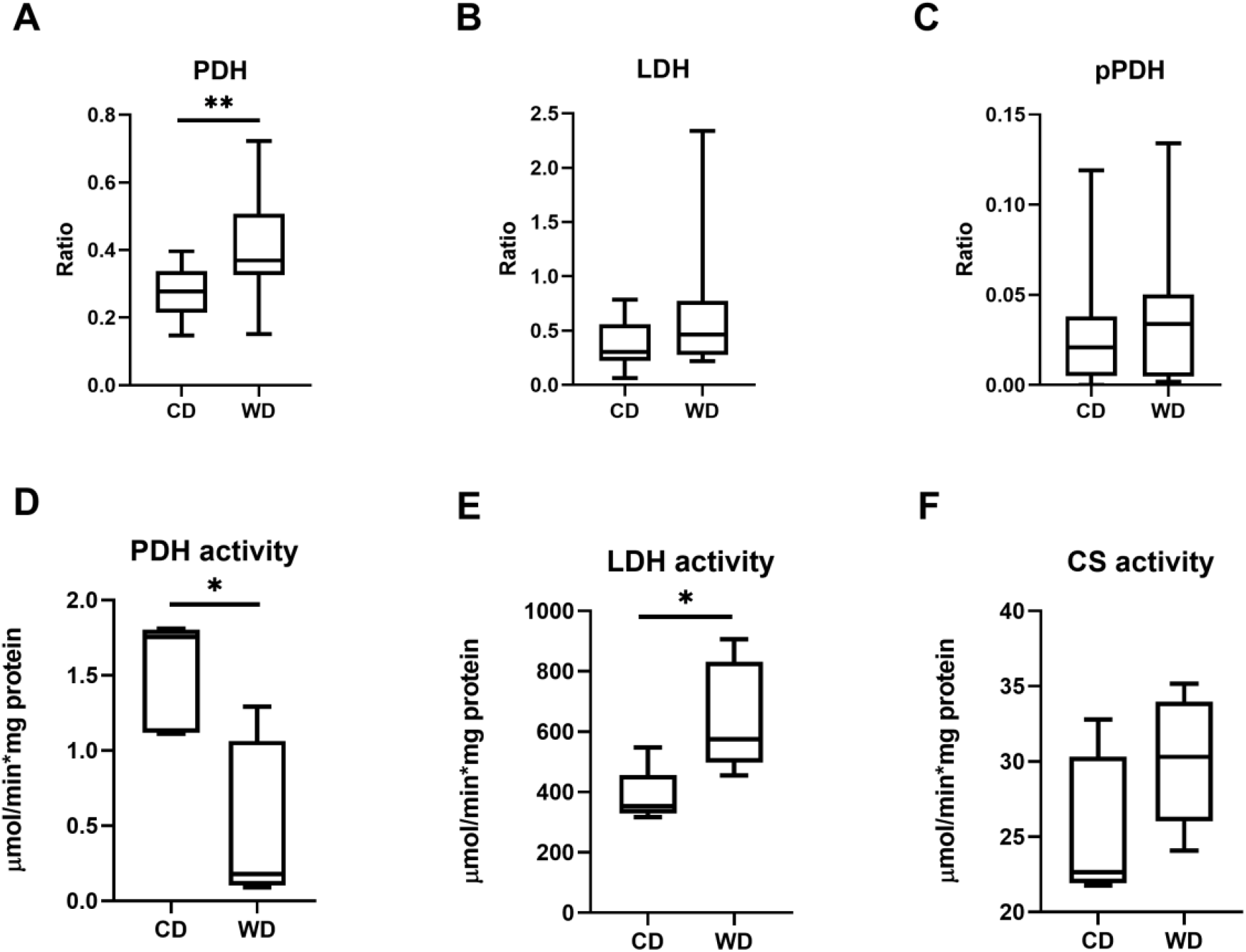
Liver enzyme protein levels for pyruvate dehydrogenase (PDH; A), lactate dehydrogenase (LDH; B), and phosphorylated PDH (pPDH; C). Enzyme activity measured from liver tissue for PDH (D), LDH (E), and citrate synthase (CS; F). * indicates p < 0.05, ** indicates p < 0.01.

### Correlations

LDH activity displayed a significantly strong positive correlation with PDFF in the liver (r = 0.8289, p = 0.0016; Figure 6A) and PDH activity displayed a significantly very strong negative correlation with liver PDFF (r = −0.8350, p = 0.0026; Figure 6B). LDH activity was also shown to have a moderate negative correlation with the lactate TTP measurement across all animals in both diet groups (r = −0.6004, p = 0.0508; Figure 6C).

**Figure 6:**
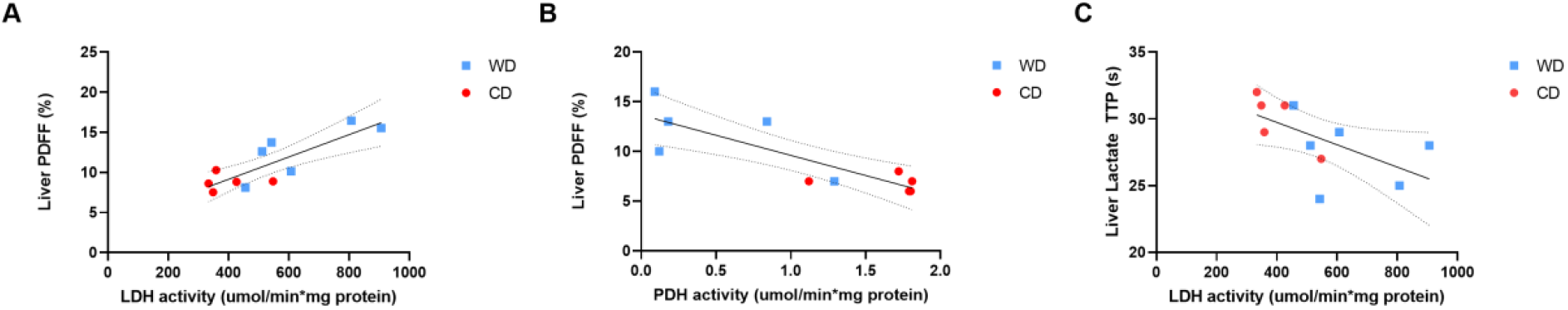
(A) Liver PDFF plotted against LDH activity displays a strong positive correlation (r = 0.8289, p = 0.0016). (B) Liver PDFF plotted against PDH activity displays a very strong negative correlation (r = −0.8350, p = 0.0026). (C) LDH enzyme activity plotted against lactate TTP displays a moderate negative correlation (r = −0.6004, p = 0.0508). Linear fit lines are shown with 95% confidence intervals.

## Discussion

### PDFF Findings Consistent with Lean NAFLD Phenotype

The increase in liver PDFF with a decline in hind limb PDFF is evidence that this model of lifelong WD consumption promotes NAFLD; differential fat deposition occurs, displaying a lean NAFLD phenotype. High fat and high fat/high sugar diets have typically been found to induce obesity in animal models (18); although lean phenotypes of NAFLD are also reported (12). PDFF values of guinea pig (19) and rat (20) livers previously reported in the literature are similar to PDFFs reported in this study for both WD and CD groups. Also of note, the liver PDFF values observed in WD animals were within the range of PDFF values (10-20%) previously reported in human patients with grade 1 and 2 steatosis (21). This pre-clinical assessment of lean NAFLD using PDFF MRI techniques similar to those that have been established in human studies (22) provides motivation for further research focused on lean NAFLD in humans which is often overlooked clinically and in the literature(23). The increased volume and weight of livers in WD-fed animals reflect fat accumulation in the liver and likely the related enlargement of hepatocytes (24). WD fed animals had significantly elevated hepatic triglyceride and cholesterol levels. The liver triglyceride concentrations in WD animals were all over 55.6 mg/g, which has been defined as a lower limit for hepatic steatosis in humans (25). Elevated liver triglyceride and cholesterol levels found in WD animals indicate that life-long exposure to WD may result in lipid overload and dysfunction in the liver (26). Related to this, elevated serum ALT was observed in WD animals, indicating liver damage (27). ALT is a reliable marker of hepatocellular injury or necrosis, suggesting that the WD group experienced liver cell damage due to diet (27). Although not statistically significant, a trend towards elevated serum ALB in WD animals was observed and is another indication of liver damage (28).

Animals in the WD group exhibited a smaller body volume, weight, and proportion of adipose tissue compared to animals in the CD group. Specifically, the decreased percentage of adipose tissue may be explained by the WD’s high fructose content, as fructose has been shown to decrease lipogenesis in rat adipose tissue (29). The overload of dietary fructose in rats has also been found to increase the release of free fatty acids from adipose tissue into the bloodstream (30), where the fatty acids may eventually be taken up by hepatocytes and stored as triglycerides in the liver (31). The loss of muscle mass has been reported in chronic liver diseases such as NAFLD (32) and we speculate this may also explain the overall decreased body weight and volume of animals fed WD in this study. While other studies have found acute high fructose diets to increase visceral adiposity (32, 33), it is speculated here that the life-long exposure to a high fructose diet during critical development stages from weaning in this study results in a reduced capacity of extrahepatic adipose tissues to synthesize and store fatty acids. Hexokinase, a hepatic enzyme responsible for fructose metabolism in adipose tissue, is adaptive to diet and may be responsible for the reduction of adipose tissue as a response to high fructose exposure (29). Similar changes of reduced body size and adiposity have been noted after exposure to high fructose diets in neonates (34) and *in utero* (35), suggesting that adaptations to an overload of fructose in the diet may be programmed at a very young age or even before birth. Previous studies in guinea pigs have reported a lean phenotype with decreased adipose tissue after life-long WD exposure with the same diet composition as the WD used in this study (19). The mechanisms of this adaptation to a high fructose diet in the adipose tissue during early development is an important topic that warrants further research.

### Altered Pyruvate Metabolism and Evidence of Damaged Livers Found in WD-fed Animals

The decreased *in vivo* lactate TTP, indicating an increased rate of lactate production, and an increased *ex vivo* LDH activity in the WD group highlight, in two techniques, a shift towards lactate production via anaerobic glycolysis in the fatty livers (36). In the current study, PDFF in the liver shows a strong correlation to LDH enzyme activity, supporting the relationship between fatty livers and the promotion of anaerobic glycolysis. Increased lactate concentration may be linked to a disturbance in liver lipid synthesis (12) and has been found in obese mice, indicating that lactate may act as a metabolic biomarker for a diet-induced fatty liver (8). WD animals show a significant increase in PDH protein levels but a significant decrease in PDH activity, suggesting inhibition of PDH in WD animals’ livers. Reports of decreased PDH activity in liver disease are associated with increased lactate as PDH normally regulates lactate production in healthy livers (37). A summary of these findings can be visualized in Figure 7. A previously published study by Lee et al. used a rat model to investigate the effects of life-long exposure to a high fat diet and found no change in lactate production, but rather an increase in aspartate and malate production in animals fed a high fat diet (38). The diet used in this study was not high in sugar, unlike the diet used in the current study, and these diet differences may explain the disparity in lactate production findings in the liver. Another relevant study used a rat model with induced obesity to investigate the effects of an acute (2-4 weeks) exposure to a high fat diet (36). This study observed increased lactate in the liver after six weeks in conjunction with NAFLD, which would be consistent with our findings of increased lactate as a result of NAFLD (36). A major difference in the acute study was the finding of increased alanine production in the liver after six weeks of the high fat diet, which is not observed in the chronic NAFLD model presented here. It may be hypothesized that the mechanisms resulting in increased liver alanine may be associated with the onset of NAFLD and become more subtle over time in a chronic model of NAFLD like the one presented in our paper. In support of this hypothesis, a study has found no evidence of elevated ALT in NAFLD patients with portal chronic inflammation, which is a marker of advanced disease state (39). Considering data from these multiple studies allows us to investigate liver function differences between acute and chronic exposure to high fat diets. Demonstrating the sensitivity of HP [1-^13^C]pyruvate MRS in measuring liver metabolism in different species and experimental conditions is an essential step in external validation for its use as an *in vivo* biomarker of liver dysfunction.

**Figure 7:**
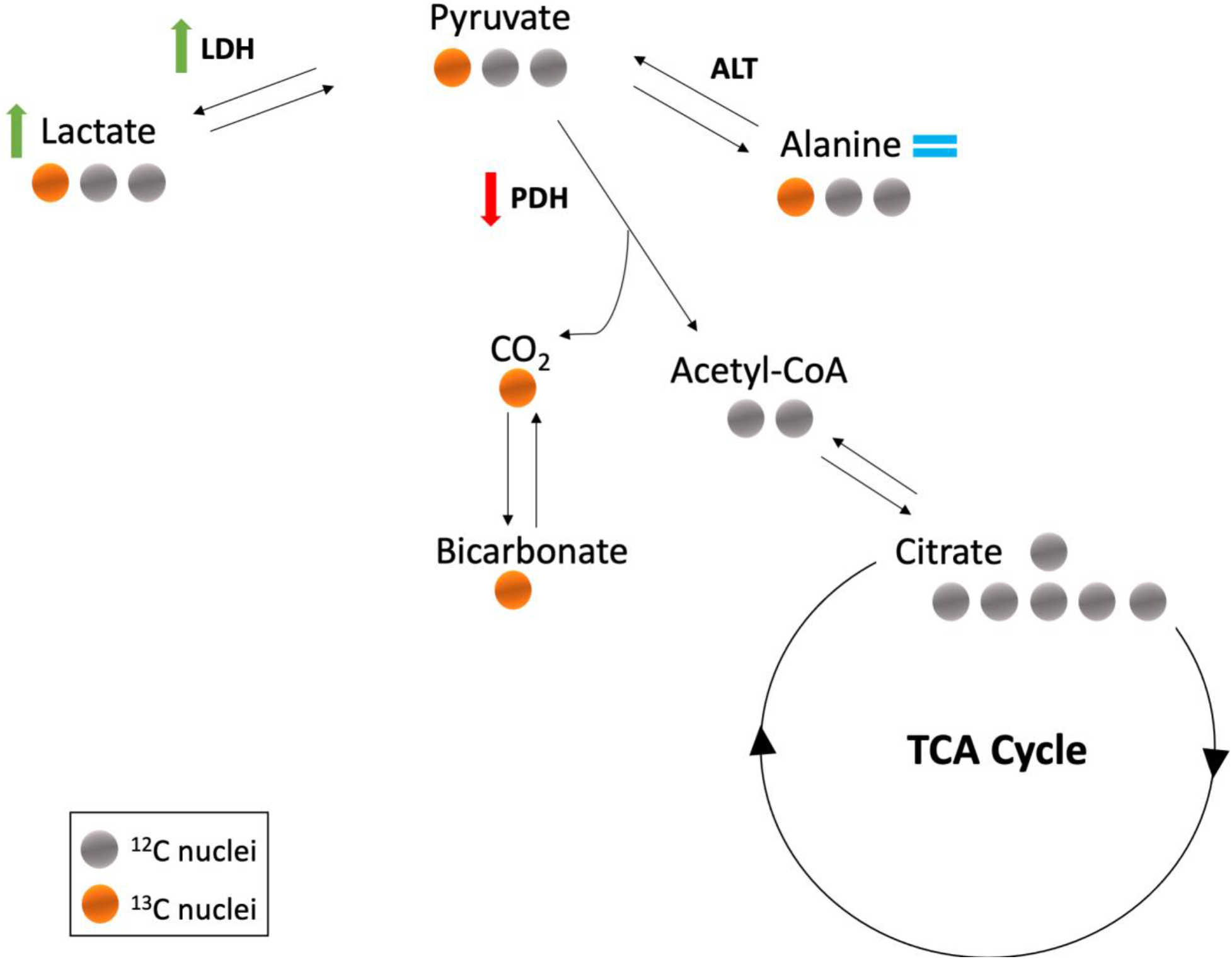
Metabolic pathways of [1-^13^C]pyruvate showing carbon nuclei as circles with ^13^C nuclei highlighted in orange. The WD group displayed an increase in lactate production and LDH activity, indicated by the green arrows, a decrease in PDH activity, indicated by the red arrow, and no change in alanine production, indicated by the blue equals sign, compared to CD animals.

### Future Work and Limitations

An approach to consider for future work may include a longitudinal study that repeats these imaging experiments in both sexes, at different time points in the guinea pigs’ development to understand the impact of sex upon these changes and also when liver metabolic changes occur in the animal’s lifespan with a life-long WD. Additionally, these imaging methods could be implemented in a study investigating whether diet reversal, exercise, or therapeutic interventions can modulate WD’s metabolic effects. Previous studies have demonstrated the benefits of dietary intervention and exercise in decreasing liver volume and fat accumulation but have not specifically looked at the impact of these interventions on hepatic pyruvate metabolism *in vivo* (40).

A technical limitation of this study is the low signal-to-noise ratio of bicarbonate in MRS experiments. Due to the low signal-to-noise ratio, there was insufficient data to measure the bicarbonate TTP. Information on the rate of bicarbonate production *in vivo* would have been valuable to correlate to PDH activity and further confirm the value of hyperpolarized ^13^C MRS as a tool to measure hepatic pyruvate[1-^13^C] labelled metabolism. Labelling the first carbon on the pyruvate molecule limits which downstream metabolites can be measured via MRS. Other experiments may label the second carbon on pyruvate instead to measure the signal from metabolites involved in the TCA cycle following the metabolism of pyruvate into acetyl-CoA. In addition, our TTP measurement precision is limited by the 1 s temporal resolution of MRS experiments. Finally, the animals’ metabolic environment may be affected by the anesthetic during MRI experiments, limiting our ability to observe homeostasis *in vivo* (41). To combat this, the animals were returned to their cages to recover in the days between the MRI examination and tissue collection. Using this method, data showing similar outcomes *in vivo* and post-MRI in collected tissue for LDH activity highlight similar readouts as evidence that changes in LDH activity may be key to a dysfunctional metabolic pathway.

## Conclusions

In summary, PDFF imaging showed an increased fat fraction in the liver, corresponding to increased triglyceride levels in the livers of male guinea pigs fed a life-long WD. Further, the application of hyperpolarized ^13^C MRS demonstrated utility in probing metabolic events in the liver that correlated with *ex vivo* liver enzyme activities. Hyperpolarized ^13^C spectroscopy results, in conjunction with altered LDH enzyme activity, highlight a shift from oxidative metabolism of pyruvate to anaerobic glycolysis and lactate production in the fatty livers of animals fed a WD. These results highlight lactate production as an indication of changes brought on by NAFLD development after chronic exposure to a WD in a guinea pig model of lean NAFLD. Hyperpolarized MRS techniques provide a non-invasive method of examining liver metabolism that correlates well with *ex vivo* findings and may be used to help determine the efficacy of interventions for NAFLD.

## Acknowledgements

We acknowledge Cheryl Vander Tuin for her assistance in ensuring the highest level of animal welfare. We also acknowledge Lindsay Morris, Simran Sethi, and Emma Coulter for their assistance with manual segmentations used to analyze the fat fraction MRI images.

